# Blood flow occlusion superimposed on submaximal knee extensions does not evoke hypoalgesia: A pilot study

**DOI:** 10.1101/2024.01.29.577836

**Authors:** Sophie-Jayne Morgan, Neil Lemay, Jenny Zhang, Neda Khaledi, Saied Jalal Aboodarda

**Affiliations:** Faculty of Kinesiology, University of Calgary, Calgary, Canada; Faculty of Physical Education and Sport Sciences, Kharazmi University, Tehran, Iran

## Abstract

Exercise-induced hypoalgesia (EIH) is a transient decrease in pain perception that can be observed following various tasks, including non-painful low-intensity and painful high-intensity exercise. The application of blood flow occlusion (BFO) can help enhance exercise adaptations while being able to exercise at a low intensity, which has important implications for clinical and rehabilitative settings. Through descending inhibitory pathways, BFO-induced pain can potentially alleviate exercise-induced pain. This study aimed to assess whether the superimposition of BFO – and its associated augmented perceived responses – during low-intensity, low-volume resistance exercise could induce hypoalgesia. Nineteen healthy adults (10 females) attended three sessions: i) no exercise (CTRL), ii) two minutes of dynamic single-leg knee extension at 10% body weight (EXER), and iii) EXER with complete occlusion applied to the upper exercising leg (OCCL). Handheld algometry-derived pain pressure threshold (PPT) of the trapezius and contralateral and ipsilateral rectus femoris muscles were measured pre- and post-exercise, and after 5 and 10 min of recovery. Perceived pain (0-10) and effort (6-20) were also rated after exercise. Although pain and effort were augmented in the OCCL condition (Pain: 6±2; Effort: 14±3) compared to CTRL (Pain: 2±2, *p<0*.*001*; Effort 9±2, *p=0*.*017*), PPT of all muscles did not change across time nor between any conditions. Therefore, the low-intensity, low-volume resistance exercise prescribed in the present study was insufficient to evoke EIH even with the application of BFO-induced pain.

## INTRODUCTION

Exercise-induced hypoalgesia (EIH) is a transient reduction in pain sensitivity to a noxious chemical, mechanical, or thermal stimulus that is often induced during moderately painful exercise (Koltyn, 2000). One of the proposed mechanisms contributing to EIH is the activation of group III/IV afferent nociceptors which transmit mechanical and metabolic perturbation signals from exercising muscles to the central nervous system during high-intensity exercise (Mense *et al*., 2001; Naugle *et al*., 2012). Integration of these signals within the brain and the consequential perceived pain is suggested to facilitate the release of endogenous opioids and activate descending pain inhibitory pathways contributing to EIH (Nahman-Averbuch *et al*., 2014; Jones *et al*., 2017). However, observing EIH in both exercised and remote, nonexercised muscles suggests that a combination of central and peripheral pain inhibition pathways is involved in this phenomenon (Naugle *et al*., 2012). In this context, the central mechanisms are primarily attributed to the release of endogenous opioids (e.g., beta-endorphins) and activation of descending pain inhibitory pathways (Janal *et al*., 1984; Olausson *et al*., 1986; Koltyn, 2000; Naugle *et al*., 2012), whereas the peripheral mechanisms are linked to the release of biochemical substances having local analgesic effects such as catecholamines and endocannabinoids in peripheral neurons (Jones *et al*., 2017).

Prior studies have suggested that performing high-intensity (Hoffman *et al*., 2004; Naugle *et al*., 2014) or high-volume exercise protocols (Krüger *et al*., 2016; Micalos & Arendt-Nielsen, 2016; Peterson *et al*., 2019) can induce hypoalgesic responses via augmented activation of group III/IV afferents. However, prescribing these types of exercise protocols is often impractical individuals experiencing chronic muscle weakness (due to injury or surgery) or musculoskeletal disorders (e.g., osteoarthritis or chronic fatigue syndrome) (Hughes *et al*., 2017; Wernbom & Aagaard, 2020). Accordingly, to increase metabolic stress and evoke EIH response, some investigators have employed blood flow occlusion (BFO) superimposed on low-intensity exercise (Suga *et al*., 2009; Ellingson *et al*., 2014; Hughes & Patterson, 2020; Hughes *et al*., 2021; Song *et al*., 2023). In this approach, the use of a pneumatic cuff at the proximal site of the peripheral limbs restricts total venous outflow and partial or full arterial inflow, which can augment metabolic stress (Thomas *et al*., 2018; Hammer *et al*., 2020) and perceived pain (Hollander *et al*., 2010) within the exercised muscles. In this scenario, activation of group III/IV afferents is postulated to increase opioid and endocannabinoid responses inducing EIH (Hughes & Patterson, 2020).

Some prior studies have observed EIH during BFO combined with continuous or multi-bout low-intensity exercises (Hughes & Patterson, 2020; Song *et al*., 2023). For instance, cycling exercise for 20 min at 40% maximal oxygen uptake (VO_2max_) superimposed with BFO resulted in EIH that was undetected during low-intensity cycling alone (Hughes *et al*., 2021). Similarly, Hughes & Patterson (2020) found the same results following 4 sets (30, 15, 15, 15 repetitions, respectively) of unilateral leg press at 30% maximal voluntary contraction (MVC) superimposed with BFO. On the contrary, Song and colleagues (2022) observed EIH after 4 sets of submaximal unilateral leg press without the addition of BFO; these investigators concluded that although BFO *could* increase the perceived volume of exercise by reducing exercise tolerance (Broxterman *et al*., 2015; Zhang *et al*., 2023; McClean *et al*., 2023), an occlusion application on its own was insufficient to induce EIH. Regardless of divergence in exercise protocols, no prior work, to our knowledge, has examined whether performing BFO during a single bout of low-intensity dynamic exercise can induce EIH. This question is critical as low-intensity, low-volume resistance training has a significant implication for the development of muscle strength in a wide range of musculoskeletal conditions.

Thus, the present study aimed to explore the effects of a single bout, submaximal intensity dynamic resistance exercise superimposed with BFO on EIH in both exercised (local) and non-exercised (remote) muscle groups. We hypothesized that the superimposition of BFO on dynamic knee extensions would induce EIH compared to submaximal exercise alone. However, due to the application of low-intensity exercise, EIH would be observed locally and not in the remote muscles.

## METHODS

### Participants

Nineteen healthy, recreationally active individuals (10 female, age: 21 ± 2 years, height: 173.9 ± 6.9 cm, weight: 70.8 ± 13.1 kg) provided written informed consent to participate in this study. Participants were eligible if they were aged 18 to 45 and were free from cardiovascular, neurological, metabolic, and/or mood disorders that would interfere with the measured variables. Participants refrained from eating and drinking for 2 hr prior, consuming analgesics, alcohol, and caffeine 12 hr prior, and lower body vigorous exercise 24 hr prior to experimental sessions. Individuals with lower extremity injuries occurring within six months of testing or those taking any analgesic medications were excluded from the study. The procedures conducted in the study were approved by the Conjoint Health Research Ethics Board at the University of Calgary (REB 21-1860).

### Experimental Protocol and Set-Up

The experimental protocol consisted of a familiarization session followed by three testing sessions completed in a randomized fashion. In the first session, after recording anthropometric measures (height and weight), participants were familiarized with the dynamic single-leg knee extension protocol, PPT and BFO procedures, and perceptual response scales, including the rate of perceived effort and pain. In the next three visits, participants completed one of the following 2-min protocols with the exercised leg: i) seated rest (CTRL), ii) one set of 2-min isotonic single-leg knee extensions using ankle weights (EXER), and iii) EXER superimposed with complete venous and arterial BFO during the knee extension contractions (OCCL). The experimental protocol was performed on an isometric chair (Kin–Com; Chattecx Corporation, Chattanooga TN, USA) while the hips and knees were fixed at 90°. The exercised leg was randomized between participants and kept constant within participants. An adjustable ankle weight (GoFit, Tulsa OK, USA) equal to 10% of each participant’s body weight was added to the ankle of their exercised leg. Knee extensions were performed through 0-90° rotation of the knee where a metronome maintained two seconds of concentric, one second isometric, and two seconds of eccentric contraction, repeated for the duration of the exercise, for a total of 24 contractions. BFO was induced using a rapid inflator system (E20 AG101; Hokanson, Bellevue WA, USA) and an adult-sized cuff placed at the proximal end of the thigh which was inflated to 300 mmHg to ensure full blood flow occlusion of the exercised leg (Azevedo *et al*., 2022; Zhang *et al*., 2023; McClean *et al*., 2023). The BFO was initiated prior to the beginning of knee extension protocol and was released once the 2-min exercise was completed. EIH was measured via pressure pain threshold (PPT), which was assessed at four separate time points in each session including: i) before the knee extension protocols (Baseline), ii) immediately after intervention protocols (Post), iii) five minutes post-intervention (5-min), and v) 10 minutes post-intervention (10-min). Baseline included two PPT trials with a 5 min gap that were averaged per session. Sessions were separated by at least 48 hours.

## Measurements

### Perceived pain and effort

Muscle pain was assessed separately using an 11-point (0–10) visual analogue scale (Cook *et al*., 1997). For this scale, participants were asked to rate the pain experienced within the exercising legs as related to the sensation of aching and burning of the muscles (whereby 0 indicated no pain at all and 10 worst possible pain) (McCormack *et al*., 1988). Pain was assessed at Baseline, Post, 5 min, and 10 min time points. Effort was measured using Borg’s 6-20 rating of perceived effort (RPE) scale (Borg, 1982) in which participants were asked to identify which numerical value best described their current exertion level. For this scale, participants were asked to rate their effort in relation to how hard, heavy, and strenuous the exercise was (whereby 6 would indicate no effort at all and 20 maximal effort) (Marcora, 2010). Effort was assessed in the last 10 s of knee extension exercises in EXER and OCCL sessions.

### Pressure Pain Threshold (PPT)

PPT was measured using a handheld pressure algometer with a stimulation area of 1 cm^2^ (Pain Test FPX 50 Algometer; Wagner Instruments, Greenwich CT, USA). PPT was assessed over the rectus femoris of both exercised and non-exercised (contralateral) legs and the upper trapezius muscle ipsilateral to the exercised muscle. To determine the test sites, the belly of the rectus femoris was palpated and marked at the distal 1/3 distance between the greater trochanter and patella, while the trapezius was identified and marked at the center point between the 7^th^ cervical vertebrae and the acromion (Vaegter *et al*., 2014; Hakansson *et al*., 2018). The rubber tip of the handheld algometer was pressed perpendicularly into the skin at the marked locations at a constant rate (approximately 30 kPa/s) by the same investigator (Ylinen, 2007). Participants were instructed to verbally indicate when the pressure turned into pain, upon which the algometer was immediately removed, and the force induced at the initiation of pain was recorded. PPT was measured twice at each site (exercised rectus femoris, non-exercised rectus femoris, and trapezius) with each location being tested once before the second measurement was taken and approximately 30 s between each measure to avoid windup effects. The average of both values from each site was used for analysis. Practice measurements were conducted on all sites in the familiarization session until participants ensured that they could identify the noxious sensation associated to their pain threshold. The order of muscles for PPT measurement was randomized between participants.

### Statistical Analysis

Descriptive analysis is presented as mean ± standard deviation. Statistical analyses were conducted using IBM SPSS, version 25.0 (IBM Corp., Armonk, NY, USA). Shapiro-Wilk test was implemented to assess data normality. For Pain and RPE, paired t tests were used to compare the 2 conditions (EXER vs. OCCL) at Post. Since no intervention was performed during CTRL, the Pain and RPE values were not expected to change; therefore, the CTRL condition was not included in the analyses of perceptual factors. Similarly, these measures were not recorded at 5- and 10-min post-assessment in the EXER and OCCL conditions. The PPT data was not normal; accordingly, Friedman’s test was used to compare 3 conditions (CTRL, EXER, OCCL) across 4 time points (Baseline, Post, 5-min, and 10-min). Statistical significance was set at an α of <0.05.

## RESULTS

### Perceived pain and effort

Paired t tests demonstrated that Pain was greater in OCCL (6 ± 2) compared to EXER (2 ± 2; *p < 0*.*001*) at Post. Similarly, RPE was greater during OCCL (14 ± 3) than EXER (9 ± 2; *p = 0*.*017*) at Post.

### Pressure pain threshold

The Friedman’s test found no significant differences between conditions or across time (within conditions) in PPT in the ipsilateral rectus femoris (*p = 0*.*150*), contralateral rectus femoris (*p = 0*.*150*), and trapezius (*p = 0*.*135*) (Figure 1).

**Figure 1.**
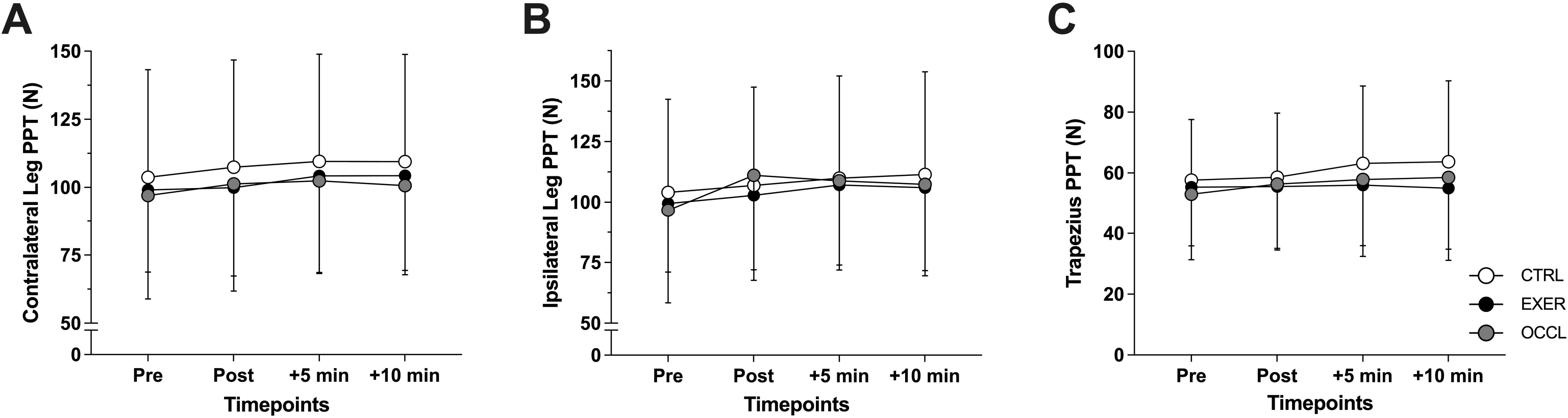
Pain pressure threshold (PPT) of the contralateral rectus femoris (A), ipsilateral rectus femoris (B), and ipsilateral trapezius (C) muscles pre-exercise, immediately post-exercise, and following 5 and 10 min of recovery in the non-exercising (CTRL; open circles), exercising (EXER; black circles), and exercising with superimposed blood flow occlusion conditions (PAIN; grey circles).

## DISCUSSION

The present study examined whether augmented perceived pain caused by BFO during a single, submaximal resistance exercise could generate EIH. We observed that while superimposition of BFO induced greater perceived pain and effort, these exacerbated perceptual responses did not induce EIH in the OCCL condition. Therefore, our results suggest that a low-volume, low-intensity unilateral resistance exercise with and without BFO is insufficient to induce a hypoalgesic response in the exercised or remote muscles.

Previous literature on EIH has indicated that high-intensity, high-volume exercise generally tends to consistently elicit EIH. For instance, protocols that employ continuous aerobic exercise at or above 70% VO_2max_ for at least 10 min tend to observe increased PPT (Hoffman *et al*., 2004; Micalos & Arendt-Nielsen, 2016; Hughes *et al*., 2021). Recent studies have also indicated that some low-intensity exercises could produce EIH resulting from possibly both systemic and local mechanisms through nitric oxide, catecholamines, endocannabinoids, and opioids released in the peripheral and central circuitry due to exercise (Jones *et al*., 2017). Kosek & Lundberg (2012) observed EIH in both the local and remote musculature after sustaining an isometric knee extension to task failure against a 1-kg weight. In another experiment, Song and colleagues (2022) observed EIH following low-intensity isometric knee extensions at 30% MVC. However, it is worth noting that in the earlier study by Kosek & Lundberg (2012), isometric contractions were sustained to task failure and in the later study by Song et al. (2022), a high-volume exercise including 4 sets of knee extensions to task failure was employed. Contrarily to these two investigations, a 3-min low-intensity, low-volume body weight wall sit also produced similar EIH results (Vaegter *et al*., 2019). Initially, these observations suggested that low-intensity, submaximal exercise could induce EIH, based on which we chose to explore the effect of a single bout, submaximal knee extension protocol in the present study. To best of our knowledge, this was the lowest exercise volume tested in the literature. Our findings did not demonstrate EIH after the selected protocol. A possible explanation for this outcome may be the total volume of exercise employed in the study being insufficient to evoke any hypoalgesic response. These findings suggest that at least a moderate volume (i.e., higher intensity and/or longer duration) of dynamic exercise may be necessary to evoke EIH. In line with this explanation, prior studies have proposed there exists a threshold for both intensity and volume of aerobic exercise that can elicit EIH (Hoffman *et al*., 2004; Naugle *et al*., 2014). However, a clear dose-response relationship between dynamic resistance exercise and EIH has not yet been established and warrants further investigation (Hoeger Bement *et al*., 2008; Vaegter *et al*., 2019).

To compensate for exercise intensity and to increase metabolic stress within exercising muscles, BFO can be superimposed upon low-intensity exercise. Hughes & Patterson (2020) examined the use of partial BFO during low-intensity exercise and compared its effects to high-intensity exercise bouts. These investigators observed that BFO combined with unilateral leg press at 30% MVC could elicit similar a EIH response as contractions at 70% MVC (Hughes & Patterson, 2020). This observation was attributed to the increased recruitment of higher threshold motor units and subsequent activation of group III/IV afferents due to the augmented perturbations in muscle milieu induced by BFO. However, this finding has been challenged by Song and colleagues (2022), who suggested that EIH was instead the result of metabolic stress prompted by reaching task failure, rather than the addition of BFO. As evidenced in the present study, despite the addition of BFO, a 2-min submaximal exercise did not elicit EIH despite increased Pain and RPE. Our findings refute the role of endogenous diffuse noxious inhibitory controls, or conditioned pain modulation (CPM), as a paradigm linked to EIH, whereby one painful stimulus can inhibit the magnitude of pain from a second, heterotopic stimulus of the same intensity via descending opioidergic and serotonergic pain inhibitory pathways (Le Bars *et al*., 1979, 1981). Although some studies have reported associations between CPM and EIH, these links are not consistently strong across the literature (Ellingson *et al*., 2014; Rice *et al*., 2019). Additionally, a caveat of the present study that should be noted is the use of handheld PPT as the only indicator of EIH. The reliability of handheld algometry can be lower compared to other pain assessment methods such as computerized algometry, dental stimulation, and cold pressor test (Janal, 1996); future studies should not only investigate the dose-response of resistance exercise coupled with BFO, but should also aim to employ multiple methods to assess pain perception to ensure greater sensitivity to EIH.

## CONCLUSION

Overall, BFO superimposed with low-intensity exercise in the present study, while did augment perceived pain and effort, did not evoke EIH. We postulate that despite the addition of BFO, the short-duration (single bout) and low-intensity (10% of body weight) of the exercise task was insufficient to activate EIH pathways, at least those measurable by handheld PPT. As no other studies, to our knowledge, have explored the effects of a unilateral single bout of low-intensity and volume exercise, this study suggests that there may be a minimum intensity of exercise needed to evoke EIH, even with BFO superimposition. Therefore, a single bout of low-intensity, unilateral dynamic exercise superimposed with BFO does not elicit EIH. These findings have implications for the use of BFO and low-intensity exercise as a pain management strategy or as prescribed exercise protocols to clinical populations unable to perform high-intensity exercise.

